# Neuropeptidome regulation after baculovirus infection. A focus on proctolin and its relevance in locomotion and digestion

**DOI:** 10.1101/2020.07.28.223016

**Authors:** Angel Llopis-Giménez, Stefano Parenti, Yue Han, Vera I.D. Ros, Salvador Herrero

## Abstract

Baculoviruses constitute a large group of invertebrate DNA viruses, predominantly infecting larvae of the insect order Lepidoptera. During a baculovirus infection, the virus spreads throughout the insect body producing a systemic infection in multiple larval tissues. Some behavioral and physiological changes in lepidopteran larvae have been described following a baculovirus infection and those changes could be connected with alterations in the host’s central nervous system (CNS). As a main component of the CNS, neuropeptides are small protein-like molecules functioning as neurohormones, neurotransmitters or neuromodulators. These peptides are involved in regulating animal physiology and behaviour and could also be targeted by baculoviruses in order to achieve host behavioural manipulations leading to increased viral fitness. In this study, we have investigated the effect of a *Spodoptera exigua multiple nucleopolyhedrovirus* (SeMNPV) infection on the neuropeptidome gene expression of *Spodoptera exigua* larval heads and brains. Expression of the gene encoding the neuropeptide proctolin was severely downregulated following infection and was chosen for further analysis. A recombinant *Autographa californica multiple nucleopolyhedrovirus* (AcMNPV) overexpressing the *S. exigua proctolin* gene was generated and used in bioassays using *S. exigua* larvae to study its influence on the viral infection. AcMNPV-proctolin infected larvae showed less locomotion activity and suffered a loss of weight compared to larvae infected with wild type AcMNPV or mock-infected larvae. These results provide additional information on the role of *proctolin* during a baculovirus infection, and offers a novel hypothesis for the molecular bases for the behavioral changes associated with a baculovirus infection.

## INTRODUCTION

Baculoviruses are a large family of entomopathogenic viruses, infecting more than 700 insect species, most of these being larvae of the order *Lepidoptera*. This specification to mainly lepidopteran insects makes them a suitable biological control agent, being safe for other insects, plants and humans (Lacey et al., 2015; Szewczyk et al., 2006). In addition, the virus is highly persistent in the environment, enhancing its potential for pest control. Nowadays, baculovirus-based biological control products are in use worldwide and novel products against different pests are under development in many countries (Moscardi et al., 2011).

One of the distinctive properties of baculoviruses is the production of two different types of virions with different functions during the infection cycle. On one hand, the occlusion-derived viruses (ODVs) contain one or multiple nucleocapsids inside a single envelope. ODVs are embedded in a proteinaceous occlusion body (OB) that causes the primary infection of the insect host. In the alkalic midgut, OBs fall apart releasing the ODVs, which bind and fuse to the midgut epithelial cells. Subsequently, the nucleocapsids enter the midgut cells, were they replicate in the nucleus. The newly formed nucleocapsids bud out of the cells, and the second type of virion is formed, budded viruses (BVs). BVs consist of a single enveloped nucleocapsid and are responsible of the secondary infection and viral spread inside the insect body. During this secondary stage of infection, baculoviruses are able to produce a systemic infection replicating in many tissues including the fat body, hypodermis, trachea, muscle and nervous system (Passarelli, 2011; Torquato et al., 2006). At the final stage of the infection, new ODVs are produced and embedded into OBs, which are released into the environment after liquefaction of the insect host. Therefore, OBs are responsible of the horizontal transmission of the virus from host to host (Granados and Lawler, 1981).

Baculovirus infections evoke behavioural changes in lepidopteran larvae, to the benefit of their own dispersion (Smirnoff, 1965). For example, *Bombyx mori* larvae infected with the B. mori nucleopolyhedrovirus (BmNPV) were found to move three to five times further than non-infected larvae (Kamita et al., 2005), a behavioural change named hyperactivity. In another example of behavioural manipulation, baculovirus infected larvae climb to the apical parts of the plant, where they eventually die and liquefy. Viral OBs are released from the corpse, spreading to the environment in what is known as tree-top disease (Goulson, 1997; Hoover et al., 2011; Van Houte et al., 2015). These behavioural changes are mediated by virus-encoded genes (Hoover et al., 2011; Kamita et al., 2005) and could be associated to changes in the host signaling pathways in the central nervous system (CNS) where part of the physiology and behaviour of the insect is controlled (Kinoshita and Homberg, 2017).

In animals, the neuropeptidergic system is composed of different neuropeptide genes (NPs) that encode small protein-like molecules used by neurons as neurohormones, neurotransmitters and neuromodulators, constituting a way of chemical communication (Van Hiel et al., 2010). These molecules are functionally related to some features of the animal’s physiology and behaviour and could be directly modulated by baculoviruses. This could lead to changes in the CNS and thus, in the insect’s physiology and behaviour, may lead to an improvement in the viral pathogenicity, virulence or its spread into the environment.

In this work we analyse how a baculovirus infection may change the neuropeptidergic system of our model organism *Spodoptera exigua* by comparative gene expression analysis in heads and brains of infected insects. Proctolin, a neuropeptide involved in the stimulation of the skeletal and visceral muscles contraction (Fiandra et al., 2010; Ormerod et al., 2016) was found to be down-regulated after an infection with S. exigua nucleopolyhedrovirus (SeMNPV). Over-expression of this neuropeptide gene and the consecutive bioassays with *S. exigua* larvae revealed its functional connection with developmental and locomotion processes. These results allow us to hypothesize about the direct or indirect modulation of proctolin by baculoviruses and its influence in important aspects of the host-pathogen interaction.

## MATERIALS AND METHODS

### Insects

Batches from the same *S. exigua* colony (SUI) were used by the two laboratories involved in this study using slightly different rearing conditions. Insects were reared at Wageningen University & Research on artificial diet at 27 °C with 50% relative humidity (RH) and a photoperiod of 14:10 h (light:dark) (Han et al., 2015). At the University of Valencia insects were reared on artificial diet at 25°C with 70 RH and a photoperiod of 16:8 h (light:dark).

### Larval infections with SeMNPV and RNA purification

Samples for the SeMNPV infection experiments were obtained from a larger study (Han et al., 2018). The larvae used for the current study were treated as follows: newly molted third instar larvae (molted and starved overnight) were fed, using droplet feeding, with a 10% sucrose solution containing SeMNPV at a 10^6^ OBs/ml concentration, which is known to kill at least 90% of infected larvae (van Houte et al., 2012). For each treatment, 30 larvae were infected by droplet feeding. As controls, 10 mock-infected larvae, droplet fed with a virus-free solution, were used per assay and none of them died due to a virus infection. Subsequently, larvae were placed individually in glass jars (120 mm high and 71 mm in diameter) containing a piece of artificial diet (aprox. 3·5 cm^3^) as described in (Han et al., 2015). Jars were placed in incubators (27 °C with 50% relative humidity and a 14:10 LD photoperiod (7 a.m. lights on, 9 p.m. lights off), with light provided from above using three luminescent tube lamps of 18 Watts each placed at a 30 cm distance above the jars. Jars were covered with transparent plastic Saran wrap containing three small holes for ventilation. Third instar mock-infected larvae (developing faster than the virus-infected larvae) were collected at 28 hours post infection (hpi). Third instar infected larvae were collected at 54 hpi. Larval heads were excised from the body of the larvae using a scalpel and total RNA was isolated from the pooled heads using the RNeasy Micro Kit (Qiagen). Three independent replicates were processed for each sample. One microgram of total RNA was used for library preparation following the TruSeq Stranded mRNA Sample Preparation Protocol (Illumina). Sequencing was performed on an Illumina HiSeq 2000 platform at Wageningen University & Research. Obtained PE-100 reads were filtered and used for gene expression analysis as described below.

### Expression analysis by RNAseq

Expression of *S. exigua* neuropeptides under SeMNPV infection were compared by mapping the trimmed reads from the different RNAseq libraries (6 replicates control and 6 replicates SeMNPV infection) to the *S. exigua* neuropeptidome (Llopis-Giménez et al., 2019). Read mapping was performed using RSEM (version 1.3.0) (Li and Dewey, 2011) and Bowtie 2 (version 2.3.4.3) software (Langmead and Salzberg, 2012). Relative abundance of each candidate gene is reported as TPM (Transcript per Million). Variability among replicates and general regulation was assessed by hierarchical clustering analysis using the heatmap.2 (gplots v.3.0.1.1) package of R software. For the statistical analysis a biological coefficient of variation (BCV) is performed with a false discovery rate (FDR) using the edgeR package (v.3.14.0) using R software.

### Expression analysis by RT-qPCR

Eighty newly molted third instar larvae (starved overnight) were fed, using droplet feeding, with a solution containing 10% sucrose solution, 1% of blue colorant Patent Blue V Sodium Salt (Honeywell) and 10^6^ OBs/ml wild type (WT) SeMNPV or a deletion mutant of SeMNPV in which the major part of the *ecdysteroid uridine 5’-diphosphate (UDP)-glucosyl transferase* (*egt*) gene was removed (Δegt-SeMNPV) virus (Han et al. 2015). In both cases, concentrations are supposed to kill at least 90% of infected larvae (Han et al. 2015). In addition, 80 control larvae were fed with the same solution containing no virus (mock infection). Subsequently, larvae were placed individually in glass jars (120 mm high and 71 mm in diameter) containing a piece of artificial diet (approx. 3-5 cm^3^) as described in (Han et al., 2015) and 2 larvae were used per jar. Jars were placed in incubators at 27 °C with 50% relative humidity and a 14:10 LD photoperiod, with light provided from above using three luminescent tube lamps of 18 Watts each placed at a 30 cm distance above the jars. Jars were covered with transparent plastic containing three small holes for ventilation. Larvae were collected at 28 hpi and 46 hpi in the case of the controls. Larvae infected with WT SeMNPV and with Δegt-SeMNPV were collected at 46 hpi (40 larvae) and 54 hpi (other 40 larvae). Six independent replicates were performed for each treatment. Larval brains were dissected in PBS and stored in 125 μl of TRIzol Reagent (Roche) at −80°C. Total RNA was purified using TRIzol reagent following the manufacturer’s instructions. One ng of each RNA sample was treated with DNAseI (ThermoFischer Scientific) following manufacturer’s protocol. Then, RNA was converted into cDNA using SuperScript II Reverse Transcriptase (ThermoFischer Scientific) following manufacturer’s recommendations and using random hexamers and oligo (dT) primers. In order to aid the nucleic acid precipitation, 10 μl of Glycogene (Roche) was used per sample.

Real-time quantitative PCR (RT-qPCR) was performed in a StepOnePlus Real-time PCR system (Applied Biosystems) using 5x HOT FIREpol Eva Green qPCR Mix Plus (ROX) (Solis Biodyne) in a total reaction volume of 20 μl. Forward and reverse primers for selected NP genes were designed using the online software tool Primer3Plus (Untergasser et al., 2007). The efficiencies for the primers were confirmed to be about 100% for the neuropeptide genes. As an endogenous control the *S. exigua ATP synthase subunit C* housekeeping gene was used in each qPCR to normalize the RNA concentration. A list of the used primers is provided in the Supplementary Table 1. The differences in expression between treatments (control and infected) were calculated using the ΔΔCt method (Livak and Schmittgen, 2001). Only expression changes statistically significant and greater than 2-fold were considered. Due to the absence of differences between the two time points, the values of the two points (46 and 54 hpi for infected samples and 28 and 46 for mock ones) were used together. Graphs and statistical analysis (unpaired *t*-test with Welch’s correction) were performed using GraphPad Prism software.

### Preparation of recombinant baculoviruses

Recombinant baculoviruses were generated using the AcMNPV Bac-to-bac Baculovirus Expression system (ThermoFischer Scientific) following the manufacturer’s instructions. The process started with constructing the donor vectors using the pFastBacDual as a backbone. A control donor vector had been previously generated in our laboratory, including the *polyhedrin* gene cloned under the *polyhedrin* promoter (to construct AcMNPV-Con). In order to generate the recombinant virus (AcMNPV-Proc) for the *Se-proctolin* gene expression, the *proctolin* gene was amplified by PCR from cDNA derived from the *S. exigua* larval brains. The used primers were F: 5’-ACTG<u>CTCGAG</u>ATGATTTCTACATTCTTGTCGGG-3’ and R: 5’-CGAT<u>CCATGG</u>TCATTCATTGAGATTACAATCT-3’ and were designed to include the XhoI and NcoI restriction sites (underlined), respectively. The *Se-proctolin* gene was cloned under the *p10* promoter, at the XhoI and NcoI restriction sites.

The donor vectors were used to transform *Escherichia coli* DH10Bac heat-shock competent cells, containing the AcMNPV bacmid, following the instructions of the manufacturer. Bacmid DNAs were recovered from *E. coli* cells and were used to transfect Sf21 cells (in Grace’s Insect medium (ThermoFischer Scientific)) using Cellfectin II reagent (ThermoFischer Scientific) following the manufacturer’s recommendations. Two independent bacmid clones from each construction were used to generate the recombinant viruses (AcMNPV-Con and AcMNPV-Proc) and these two strains per recombinant virus were used as biological replicates in the subsequent bioassays. After 5 days of incubation at 27°C, the resulting baculoviruses were collected (P0). Two more steps of baculovirus amplification were performed in Sf21 cells in order to obtain the viral stock (P1 and P2). P2 OBs were purified using a 40% sucrose gradient protocol (Caballero et al., 2001). The OBs used in the bioassays were quantified in the Neubauer chamber as OBs per millilitre (OBs/ml). The budded virus used for the growing curves were quantified by qPCR as was previously described (Martínez-Solís et al., 2017). The virus concentrations were calculated in budded virus per millilitre (BVs/ml).

The time-course replication of both types of recombinant viruses were assessed in Sf21 cells. The two independent clones of AcMNPV-Con and the two clones of AcMNPV-Proc were used to infect Sf21 cells in a 12-well plate (1 ml of cells/well). This was performed in three replicates. Each 24 hours (from 0 to 96 hours post infection (hpi), 25 μl of each well supernatant was taken for viral DNA extraction and quantification. DNA was treated with Prepman Ultra Sample Preparation Reagent (ThermoFischer Scientific) and was directly used for qPCR, using *DNA polymerase* primers and a standard curve of known amounts of viral DNA (from 10 ng to 0.001 ng). Values were represented using GraphPad Prism software.

### Insect bioassays: pathogenicity and virulence

Two bioassays were performed, one with a sublethal viral dose (5·10^5^ OBs/ml) and a second one with a lethal viral dose (1·10^7^ OBs/ml). Three treatments were used per bioassay: mock infection, infection with AcMNPV-Con and infection with AcMNPV-Proc. Thirty-two newly molted *S. exigua* third instar larvae were used per treatment and replicate. Infections were performed by droplet-feeding, in drops containing the virus solution or mQ water (mock controls), 10% sucrose and 10% Phenol Red as mentioned above. After that, larvae were individualized in trays with a piece of artificial diet and mortality was recorded every 12 hours until the end of the bioassay (286 hpi). At the end of the bioassay, mortality rates were calculated for each of the virus and the different doses. Graphs and statistical analysis (Gehan-Breslow-Wilcoxon survival curve test) were performed using GraphPad Prism software. Assays were done in three independent replicates involving different batches of insects and using two different viral clones for each of the recombinant viruses.

### Insect bioassays: digestion and locomotion

In order to analyse how *proctolin* could affect digestion processes, 32 newly molted *S. exigua* third instar larvae were used per treatment and per replicate and infected as described above. For these assays larvae were not individualized. The weight of the larvae during the bioassay was monitored every 24h from the start of the bioassay. Larvae from each treatment were weighted in groups in order to estimate average individual weight. From 72 hpi to 144 hpi, stools were taken from the bioassay trays and were weighted every 24 hours. Graphs and statistical analysis (two-way ANOVA) were performed using GraphPad Prism software. Assays were done in three independent replicates involving different batches of insects and using two independent viral clones for each of the recombinant viruses.

The possible influence of *proctolin* on larval locomotion was also analysed. Ten *S. exigua* larvae were placed in one side of a 14 cm diameter Petri dish whereas a piece of artificial diet (1.5 x 0.8 x 1 cm) was placed at the opposite side. The Petri dish was placed inside a paperboard box (30 x 22 x 22 cm). A hole in the side of the box (6 cm of diameter) was made to include a 50 W halogen artificial light (at 15 cm of distance to the Petri dish) (Figure 6A). For each replicate, larval mobility was scored by dividing the Petri dish in 10 areas of 1.3 cm each (figure 6A). At time 2’, 5’ and 10’ the number of larvae in each area was recorded and then, the interval travelled by each them was averaged and converted into cm, obtaining a mean mobility index. This bioassay was performed with the three groups of larvae: mock-infected larvae, larvae infected with AcMNPV-Con and larvae infected with AcMNPV-Proc, in a total of six replicates. Graphs and statistical analysis (two-way ANOVA) were performed using GraphPad Prism software.

### Proctolin expression in larvae heads

In order to quantify the native and recombinant proctolin in bioassayed larval heads, groups of 4 heads were taken from each group of larvae at different time points: 96 hpi and 120 hpi from the mortality assay; and 120 hpi and 144 hpi from the digestion and locomotion assays. The heads were stored in 200 μl of TRIzol and RNA extraction was performed following maker’s instructions. 500 ng of each RNA were treated with DNAseI and converted into cDNA as described before.

RT-qPCR analysis was performed as previously described. Different primers were used in the analysis: AcMNPV DNApol, for quantifying viral presence in the samples; *S. exigua* proctolin, for quantifying the native proctolin; and AcMNPV proctolin, for recombinant proctolin detection (the reverse primer hybridizes with a 3’ region in the baculovirus expression vector). A list of the used primers is provided in the Supplementary Table 1. The cycle threshold (Ct) values from each sample was used to know the relative expression level of the different genes through the 2^-Ct-Proc^/2^-Ct-DNApol^ method. Graphs were performed using GraphPad Prism software.

## RESULTS

### Neuropeptide expression in larval heads after SeMNPV infection

Effect of viral infection on the expression of the *S. exigua* neuropeptidome was initially assessed in larval heads. The overall analysis did not reveal major difference in global regulation of neuropeptides expression between control (mock-infected) and virus-infected samples as the clustering analysis of the expression values could not group samples by their treatment (figure 1). Nevertheless, detailed analysis of each one of the different genes revealed nine neuropeptides differentially expressed following baculovirus infection. From the 73 analysed neuropeptides, two of them appear upregulated: insulin-like peptide 1 (*ILP1*) (8.3-fold change) and insulin-like peptide 2 (*ILP2*) (2.3-fold change). Seven neuropeptide genes were downregulated: ecdysis triggering hormone (*ETH*) (6.3-fold change), ion transport peptide (*ITP*) (6-fold change), PBAN-DH (1,7-fold change), eclosion hormone (*EH*) (5.2-fold change), SIFamide (*SIF*) (4-fold change), leucokinin (*LK*) (2.1-fold change) and prothoracicotropic hormone (*PTTH*) (2.3-fold change).

**Figure 1.**
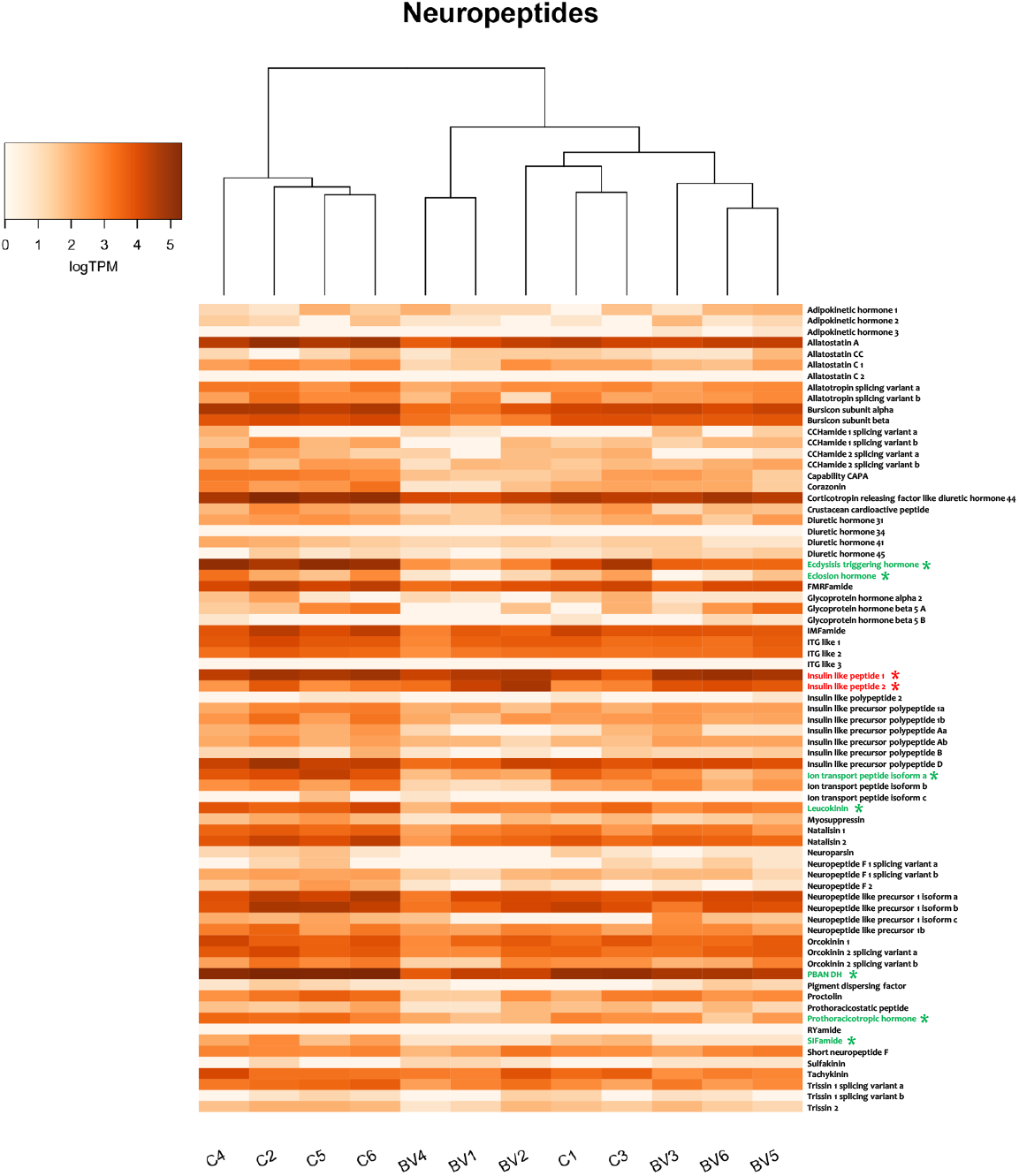
Heat-plot of the relative expression of neuropeptides in the head of *S. exigua* larva infected by SeMNPV. Estimation of abundance values was determined by read mapping. Colour plots represent values of TPM logarithm. Light orange colours indicate low expression and dark orange ones indicate high expression. C: Control sample. BV: Baculovirus-infected sample. Asterisks indicate statistically significant differences between non-infected and infected samples (P<0.05). Red and green labelled genes are indicative of up-regulation or down-regulation, respectively.

### Neuropeptide expression in larval brains after SeMNPV infection

Larval brain accounts for a small fraction of the total head mass and constitutes the main tissue where most of NPs are expressed (Fónagy, 2014; Nässel and Homberg, 2006). Differential expression analysis of selected NPs in the brain of virus-infected insects was performed through RT-qPCR. 21 NPs were selected according to the previous expression results and their potential role in behavioural. Experiments were conducted with wild type (WT) SeMNPV and Δegt-SeMNPV infected larvae to check whether the lack of the *egt* gene could influence the NPs expression. The presence of viral activity in the brain tissues, measured as the expression of the viral *DNApol* gene, was confirmed and found similar for both viruses (WT and Δegt) (Figure 2A and B). The overall transcription pattern of the selected NPs was similar for both viral infections. Independently of the viral genotype, most of the analysed genes were not showing important changes in gene expression following viral infection. From the differentially expressed genes, *proctolin* appears to be significantly down-regulated for the two viral infections. *Proctolin* expression was 5.6- and 5.4-fold down regulated after infection with WT SeMNPV and the Δegt-SeMNPV, respectively.

**Figure 2.**
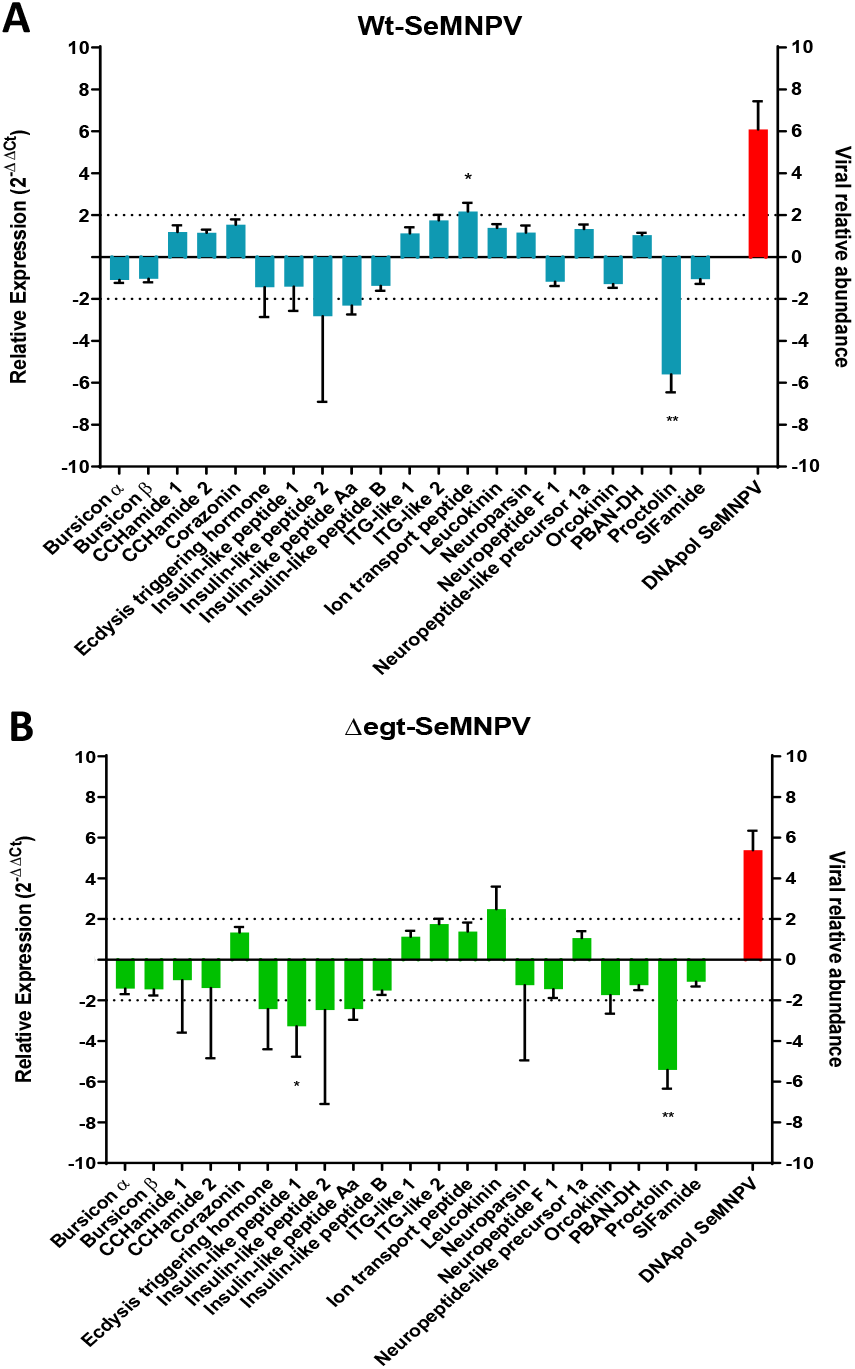
Differential expression analysis in *S. exigua* larval brains after infection with SeMNPV. A) WT-SeMNPV and B) Δegt-SeMNPV. RT-qPCR assays were performed using gene specific primer pairs. Asterisks indicate statistically significant differences between non-infected and infected samples (P<0.05). The red column shows the expression of the SeMNPV DNApol gene.

### Effect of proctolin expression on larval physiology and behaviour

Motivated by the consistent and reproducible downregulation of *proctolin* in larval brains after viral infection, additional studies were conducted to dissect the influence of *proctolin* on the physiology and behaviour of *S. exigua* larvae during a viral infection. For that purpose, a recombinant baculovirus constitutively expressing the *Se-proctolin* gene (AcMNPV-Proc) was constructed. AcMNPV-Proc replicates in Sf21 cells similarly to the counterpart control (AcMNPV-Con) (Figure 3A). In addition, the expression of recombinant and native *proctolin* in the brain of AcMNPV-infected larvae was confirmed using specific primers (Figure 3B). Recombinant proctolin was highly expressed in the brain of larval infected with the AcMNPV-Proc virus.

**Figure 3.**
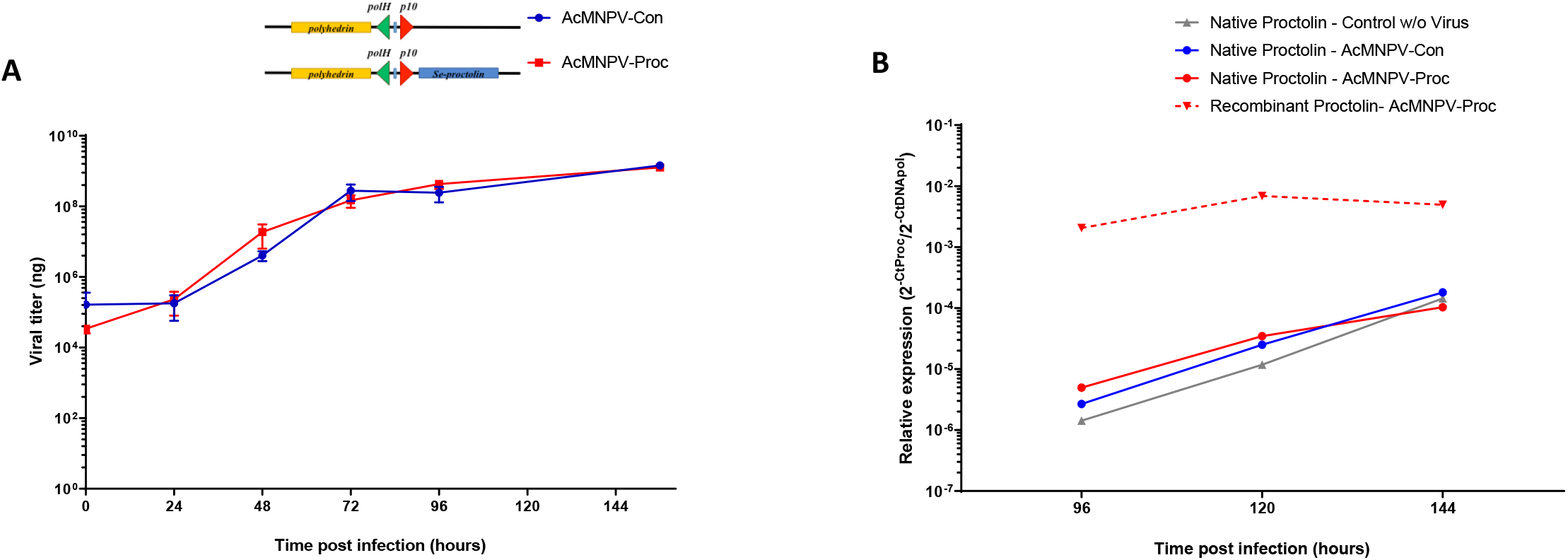
Viral growth and *proctolin* expression in AcMNPV viruses. A) Growth curves of AcMNPV-Con and AcMNPV-Proc in Sf21 cell culture. The image above the graph shows a schematic representation of the viral constructs. B) Relative expression of native and recombinant SeProctolin in bioassayed *S. exigua* larval heads. RT-qPCR assays were performed using gene specific primer pairs. Native proctolin in the non-infected samples appears in grey, native proctolin in AcMNPV-Con samples appears in blue, native proctolin in AcMNPV-Proc samples appears in continuous red and recombinant proctolin in AcMNPV-Proc samples appears in dotted red.

Effect of the over-expression of *proctolin* in the baculoviral pathogenicity and virulence was assessed by bioassaying AcMNPV constructs at sublethal (5·10^5^ OBs/ml) and lethal (1·10^7^ OBs/ml) doses. At the sublethal dose, AcMNPV-Proc was more pathogenic (45% mortality) that its control virus (25% mortality) (Figure 4A). However, at the lethal doses, no effect of *proctolin* expression on viral activity was observed and similar pathogenicity and virulence was found for both viruses (Figure 4B).

**Figure 4.**
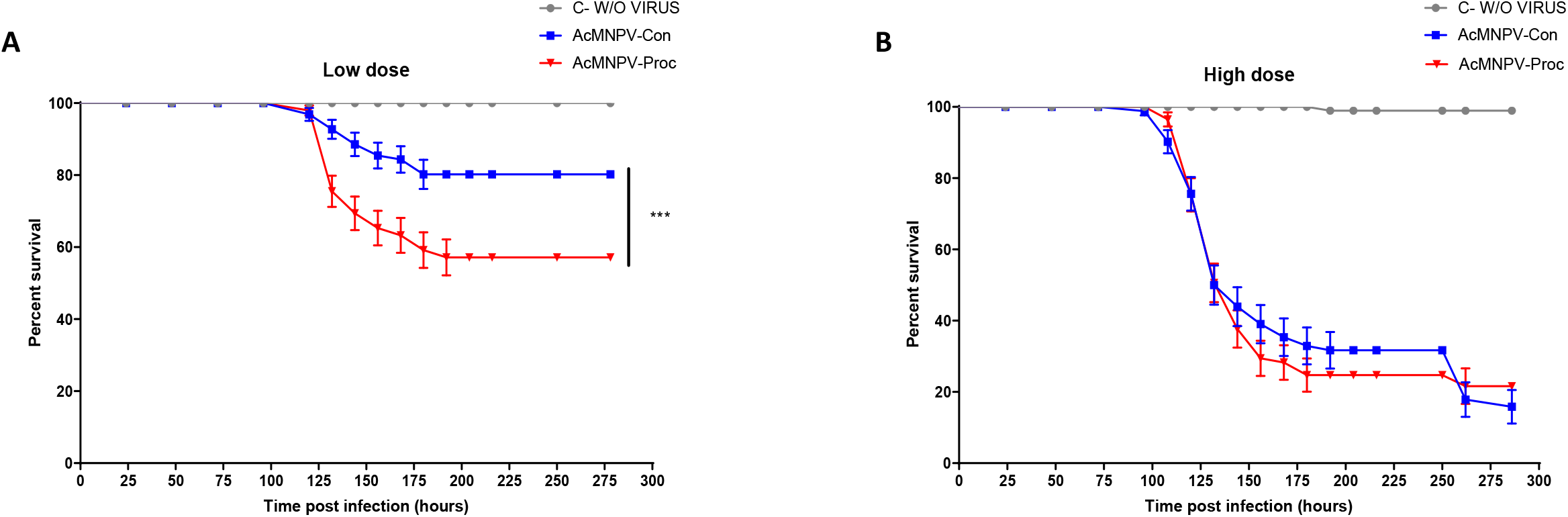
Pathogenicity of the SeProctolin-expressing viruses. Larval mortality was assessed at two viral concentrations. A) Mortality at sublethal viral dose. B) Mortality at lethal viral dose. Asterisks indicate statistically significant differences between AcMNPV-Con and AcMNPV-Proc samples (P>0.05).

The lethal viral dose was then selected for further experiments to assess the effect of *proctolin* on physiological parameters previously associated to the activity of this peptide in insects (Fiandra et al., 2010; Ormerod et al., 2016). Specifically, influence of virus-expressed *proctolin* was studied on the larval growth, digestion rates, and locomotion.

*Proctolin* over-expression during the baculovirus infection was found to reduce the larval growth as reflected in a reduction in the larval weight and larval stools weights (Figures 5A and B). Differences between larvae infected with AcMNPV-Con and AcMNPV-Proc started to appear at 96 hpi, reaching out to a reduction of about 25% of the larval weight in AcMNPV-Proc infected larvae when compared to the non-infected or AcMNPV-Con-infected larvae at 144 hpi (p-value <0.001). Similarly, larval stool weight was also reduced in the larvae infected with the *proctolin*-overexpressing virus. When compared to larvae infected with AcMNPV-Con, the stool weight of larvae infected with AcMNPV-Proc was reduced in about a 50% (p-value <0.001) at 144 hpi. These results suggest that *proctolin* over-expression affects, directly or indirectly, to the insect digestion and food intake, and consequently to the larval development.

**Figure 5.**
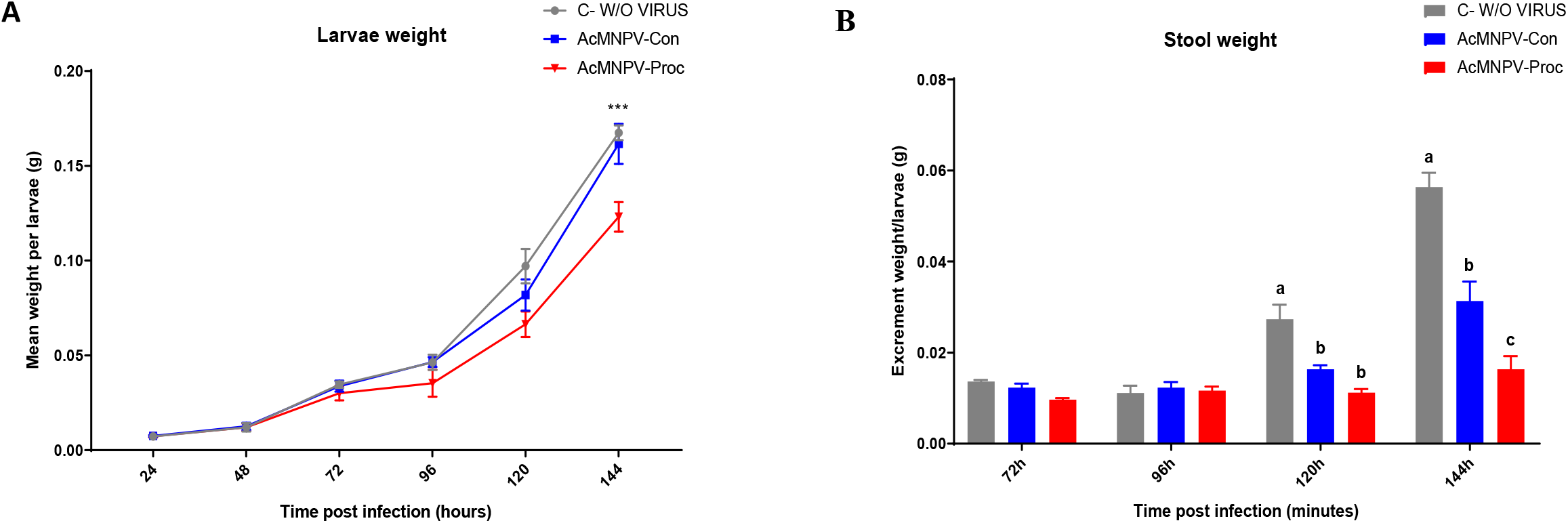
Effect of SeProctolin in larval growing and digestion. A) Mean larvae weight of *S. exigua* infected larvae at different time points. Asterisks indicate statistically significant differences between AcMNPV-Proc and AcMNPV-Con (P>0.05). B) Mean stool weight of *S. exigua* infected larvae at different time points. Letters indicate statistically significant differences between the sample groups (P>0.05).

Regulation of larval locomotion has also been associated to the action of proctolin (Ormerod et al., 2016). Our locomotion assay revealed that *proctolin* over-expression reduced, directly or indirectly, the locomotion activity of the larvae. At 144 hpi larvae infected with the AcMNPV-Proc virus have less locomotion activity, compared to the controls (Figure 6B). When compared to the non-infected and the AcMNPV-Con infected larvae, the infection with AcMNPV-Proc showed a significant reduction in about 70% in the locomotion value (p-value <0.001).

**Figure 6.**
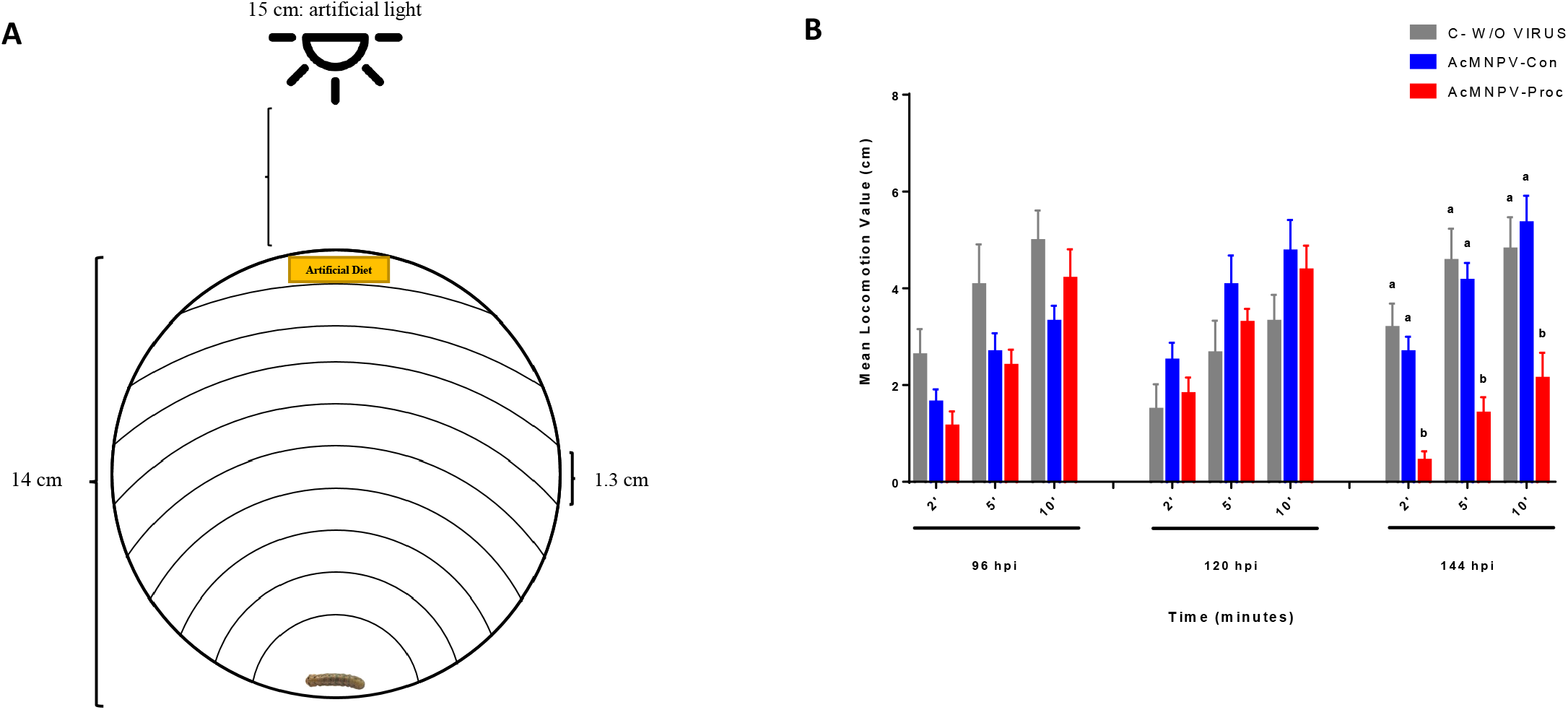
Effect of SeProctolin in larval locomotion. A) Schematic representation of the behaviour assay set up (see material and methods). B) Mean locomotion value of *S. exigua* mock and infected larvae. Letters indicate statistically significant differences between the sample groups (P>0.05).

### Proctolin effects on expression of other NPs

Effect of *proctolin* over-expression on the regulation of other NPs in the brains of the larvae was also studied to check if the observed effects in development and locomotion could be due to the unique action of *proctolin* or to a dis-regulation of the neuropeptidergic system after *proctolin* over-expression. For that, expression of about 20 NPs was compared by RT-qPCR in the brain of AcMNPV-Con and AcMNPV-Proc infected larvae (144 hpi). Genes were selected because of the already described function and their role connected to developmental or locomotional aspects of the physiology. None of the analysed genes was found to be differentially regulated by the *proctolin* overexpression (Figure 7) suggesting that observed changes in digestion and locomotion were mainly attributed to the direct action of *proctolin*.

**Figure 7.**
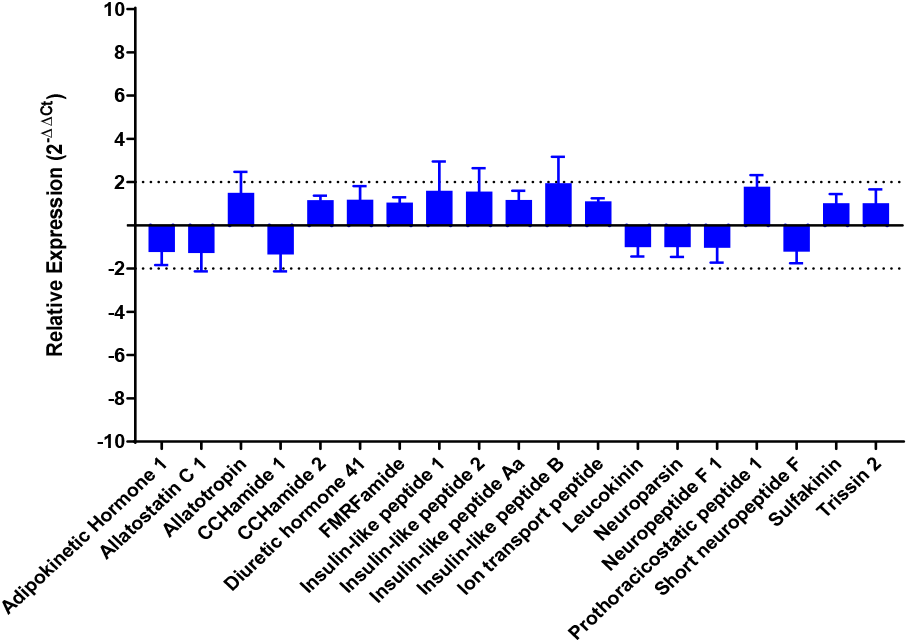
Differential expression analysis in *S. exigua* larva head samples under AcMNPV-Proc infection. RT-qPCR assays were performed using gene specific primer pairs.

## DISCUSSION

Baculoviruses interact with their host in many ways and have developed multiple strategies to increase their fitness and dispersion through the alteration of the physiology an behavior of the host (Cheng and Lynn, 2009; Kong et al., 2018). The neuropeptidergic system in insects is composed of signalling molecules playing a role in the chemical communication between cells (Elphick et al., 2018; Schoofs et al., 2017). Due to its regulatory function of the insect’s physiology and behaviour (Bendena, 2010; Schoofs et al., 2017), understanding their regulation after baculovirus infection would contribute to understanding some of the physiological and behavioural changes that have been associated to baculovirus infections. To study this, the expression levels of the genes encoding the *S. exigua* neuropeptidome (Llopis-Giménez et al., 2019) following SeMNPV infection was initially analysed in the whole head of the larvae. Although no clear pattern of regulation of the neuropeptidergic system was detected, few NPs were found to be differentially expressed during infection. Two insulin-like peptides were the only upregulated genes. They are functionally related to the larval development and belong to a wide family of genes (Wu and Brown, 2006). Seven neuropeptides were downregulated although no major connection was found between them except for the Ecdysis Triggering Hormone and Eclosion Hormone that are related to the ecdysis (molting) process. Downregulation of these two genes could produce a delay in the ecdysis, a phenotype already connected with the baculovirus infective process (O’Reilly and Lois K. Miller, 1991).

More specific analyses of the changes in the larval brain, revealed a different set of regulated NPs when compared to the changes observed using the complete larval heads. Although neuropeptides are also expressed in other organs including endocrine glands in the intestinal tract and in muscles, the brain is the main organ in which they are expressed (Fónagy, 2014), representing only a small portion of the head. RNA-seq analysis from composite tissues as larval head make it more difficult to interpret positive signals for genes with variable patterns of local expression (Johnson et al., 2013) and could also mask changes in the CNS. Accordingly, although changes observed using the whole head have provided information on some NPs affected during the baculovirus infections, those changes observed in the brain are more likely to be involved in the behavioural changes produced during a baculovirus infection.

Gene expression analysis of larval brain revealed a clear down-regulation of *proctolin* following an SeMNPV infection. Down-regulation of *proctolin* was consistent among the different replicates and similar for the two viral strains used in the study. Proctolin was first discovered in *Periplaneta americana* as a five amino acid cyclic peptide that inhibited the contractions of the hindgut. The presence of proctolin and its function as a myotropic neurotransmitter and neuromodulator has been additionally confirmed in the dipteran *D. melanogaster* (Anderson et al., 1988; Isaac et al., 2004), in the coleopteran *Tenebrio molitor* (Breidbach and Dircksen, 1989) and in the orthopteran *Locusta migratoria* (Clark et al., 2006). In the hemipteran *Rhodnius prolixus, proctolin* stimulates the contractions in the midgut and the hindgut (Orchard et al., 2011). In lepidopteran insects, proctolin was first studied in *Pieris rapae* in which it was found to have a variable activity in the hindgut contractions (Walker and Bloomquist, 1999). In *Bombyx mori*, proctolin was found to increase the contractile activity in the larval hindgut at high concentrations. Moreover, when it was orally administered, a significant reduction in the conversion of ingested and digested food was observed, confirming its role in growth and development (Fiandra et al., 2010). In the same study, the authors also reported the reduction of *Spodoptera littoralis* larval growth by a 20% after proctolin treatment (Fiandra et al., 2010). To gain information on the role of proctolin during baculovirus infection, a gain-of-function strategy was chosen and an AcMNPV baculovirus constitutively expressing *Se-proctolin* was constructed. Infections with AcMNPV-Proc showed an important reduction in larval growth and digestion (Figure 5) when compared with the control AcMNPV (not expressing *Se-proctolin*). Those experiments confirm the functionality of the recombinantly expressed precursor and reveal similar functions for this NP in *S. exigua* larvae. In previous studies on *S. exigua*, proctolin was found to be expressed in the larval head (neurons of the CNS) and in the midgut (endocrine cells) (Llopis-Giménez et al., 2019), confirming its classification as brain-gut peptide and coinciding with previous descriptions in other organisms as *D. melanogaster* (Llopis-Giménez et al., 2019; Taylor et al., 2004). In our approach, expression of SeProctolin was not restricted to the larval brain and whether the observed effect with the SeProctolin overexpression was due to the direct effect of the NP on the larval gut, as an endocrine messenger, or a combination of both remains to be elucidated. In other systems, as in *B. mori* larvae, proctolin was confirmed to trespass the midgut epithelium and reach its haemocoelic receptors, confirming its direct and myotropic effect in the increase of midgut contractions and its consequences in the food absorption (Fiandra et al., 2010, 2009). In *D. melanogaster*, muscle contractions were absent when they knocked down the proctolin receptor. They also observed that proctolin was acting downstream of the postsynaptic depolarization regulating the calcium influx (Ormerod et al., 2016). This has also been confirmed in other organisms as in the crustacean *Isodea baltica*, where proctolin directly decreases the membrane conductance, increasing the excitability of the gut muscles (Erxleben et al., 1995).

In addition, proctolin seems to also act as a cotransmitter enhancing the neuromuscular transmission and the skeletal muscle contractions (Orchard et al., 1989). In fact, proctolin was suggested to be associated with a slow motor function affecting the locomotion activity of the insects although this effect was never directly showed (Lange and Orchard, 2006; O’Shea and Bishop, 1982; Witten and O’Shea, 1985). After confirming the functionally of the SeProctolin expressed in AcMNPV we have also explored the effect of the SeProctolin over-expression on the locomotion of the larvae, a parameter that has been associated to the behavioural changes that occurs during the baculovirus infections (Goulson, 1997; Kamita et al., 2005). To study the locomotion activity of the larvae a locomotion assay was developed. This new method resulted in an efficient set up able to detect differences between the treatments linking for first time the proctolin activity with a reduction of the larval locomotion.

Baculovirus infection produces an enhanced locomotion activity (ELA), also known as hyperactivity, in infected caterpillars. In addition, baculoviruses cause caterpillars to climb to the upper parts of the plant, in a phenotype known as *Wipfelkrankheit* or tree-top disease. Hyperactivity has been related to the viral *ptp* gene, that encodes the protein tyrosine phosphatase (PTP) and tree-top disease has been related to the viral *egt* gene that encodes the ecdysosteroid UDP-glucosyltransferase (EGT) (Hoover et al., 2011; Kamita et al., 2005). PTP seems to be involved in the wandering-like ELA enhanced by light and related with a positive phototropism (Kamita et al., 2005). EGT is involved in the life extension of the larvae and the vertical climbing behavior (Han et al., 2015; Hoover et al., 2011), although an effect of EGT on tree-top disease was not observed for all virus-host combinations (Ros et al. 2015). This two genes favour two different behavioural changes that could be complementary in many cases. Although these two genes have been connected to these behavioural changes, the underlying mechanisms may be more complex involving modifications in the CNS, where the main behavioral aspects of the host are controlled (Gasque et al., 2019; Katsuma et al., 2012). It is tempting to hypothesize that the down-regulation of *SeProctolin* as a consequence of the baculovirus infection in the larval brain could be connected to hyperactivity, as a decrease in the expression of this neuropeptide may produce an increase in the locomotion activity in the larvae. Nevertheless, down-regulation of *proctolin* was produced under the infection of WT SeMNPV as well as the Δegt-SeMNPV. That implies that the presence of the *egt* gene is not associated with the regulation of the *SeProctolin* gene expression nor with locomotion changes mediated by this neuropeptide. The observed changes in the larval digestion and development associated to the *SeProctolin* over-expression would be also independent from the *egt* gene, although functionally this baculovirus-expressed gene has been related with the upkeep of the larvae in an active feeding state, producing bigger larvae and increasing the yield of viral progeny. We hypothesize that the decrease in *SeProctolin* expression observed in larval brain would complement the *egt* effects, producing bigger larvae possibly due to a decrease in the midgut contraction rates.

Whether the reduction in *proctolin* expression after SeMNPV infection is the result of the direct effect of the virus on the host, a defence response of the host, or just a side effect of the physiological changes associated to the infection, remain to be elucidated. Down-regulation of host genes to increase viral fitness or evade the host’s immune response by viral encoded miRNAs has been described for several baculovirus species (Kharbanda et al., 2015; Singh et al., 2010; Tang et al., 2019). At present, our analysis of SeMNPV-expressed miRNAs has not revealed miRNAs targeting to *SeProctolin* (data not shown).

*SeProctolin* downregulation has been described in this work as a consequence of the SeMNPV infection in *S. exigua* larval brains. Using a *SeProctolin* over-expression approach, we have confirmed its role in regulating physiology aspects as growing and locomotion. Although additional research will be needed to unveil the regulation mechanism of *proctolin* by SeMNPV, these results allows us to hypothesize about the role of this NP in the baculovirus-host interaction and the previously described behavioral changes produced by baculoviruses that contribute to viral dissemination.

## Acknowledgements

This study was supported by the Spanish Ministry of Science, Innovation and Universities (No. AGL2014-57752-C2-2R and RTI2018-094350-B-C32). ALG was recipient of a PhD grant from the Spanish Ministry (grant BES-2015-071369). ALG was also recipient of an EMBO Short-Term Fellowship (grant 7199) for a two-month research period at Wageningen University & Research (the Netherlands). We thank Rosa María González-Martínez, Oscar Marín-Vázquez, Els Roode and the late Hanke Bloksma for their excellent help with insect rearing and laboratory management.

## REFERENCES

Anderson, M.S., Halpern, M.E., Keshishian, H., 1988. Identification of the neuropeptide transmitter proctolin in Drosophila larvae: Characterization of muscle fiber-specific neuromuscular endings. J. Neurosci. 8, 242–255. https://doi.org/10.1523/jneurosci.08-01-00242.1988

Bendena, W.G., 2010. Neuropeptide physiology in insects. Adv. Exp. Med. Biol. 692, 166–191. https://doi.org/10.1007/978-1-4419-6902-6_9

Breidbach, O., Dircksen, H., 1989. Proctolin-immunoreactive neurons persist during metamorphosis of an insect: A developmental study of the ventral nerve cord of Tenebrio molitor (Coleoptera). Cell Tissue Res. 257, 217–225. https://doi.org/10.1007/BF00221653

Caballero, P., Cisneros, J., Claus, J.D., Cherry, A., Del Rincón, M.C., Ghiringhelli, D., Ibarra, J.E., López-Ferber, M., Luque, T., Martínez, A.M., Moscardi, F., Muñoz, D., Murillo, R., Romanowski, V., Ruiz de Escudero, I., Sciocco de Cap, A., Sosa-Gómez, D.R., Osuna, E.V., Vasconcelos, S.D., Vilaplana, L., Williams, T., 2001. Los baculovirus y sus aplicaciones como bioinsecticidas en el control biológico de plagas. Universidad Pública de Navarra - Phytoma España, Navarra.

Cheng, X.W., Lynn, D.E., 2009. Baculovirus Interactions In Vitro and In Vivo. Adv. Appl. Microbiol. 68, 217–239. https://doi.org/10.1016/S0065-2164(09)01205-2

Clark, L., Agricola, H.J., Lange, A.B., 2006. Proctolin-like immunoreactivity in the central and peripheral nervous systems of the locust, Locusta migratoria. Peptides 27, 549–558. https://doi.org/10.1016/j.peptides.2005.06.027

Elphick, M.R., Mirabeau, O., Larhammar, D., 2018. Evolution of neuropeptide signalling systems. J. Exp. Biol. 221. https://doi.org/10.1242/jeb.193342

Erxleben, C.F.J., DeSantis, A., Rathmayer, W., 1995. Effects of proctolin on contractions, membrane resistance, and non-voltage-dependent sarcolemmal ion channels in crustacean muscle fibers. J. Neurosci. 15, 4356–4369. https://doi.org/10.1523/jneurosci.15-06-04356.1995

Fiandra, L., Casartelli, M., Cermenati, G., Burlini, N., Giordana, B., 2009. The intestinal barrier in lepidopteran larvae: Permeability of the peritrophic membrane and of the midgut epithelium to two biologically active peptides. J. Insect Physiol. 55, 10–18. https://doi.org/10.1016/j.jinsphys.2008.09.005

Fiandra, L., Casartelli, M., Diamante, B., Giordana, B., 2010. Proctolin affects gut functions in lepidopteran larvae. J. Appl. Entomol. 134, 745–753. https://doi.org/10.1111/j.1439-0418.2009.01501.x

Fónagy, A., 2014. Insect Neuropeptides and their Potential Application for Pest Control Insect Neuropeptides and their Potential. Acta Phytopathol. Entomol. Hungarica 41 (1–2), 137–152. https://doi.org/10.1556/APhyt.41.2006.1-2.13

Gasque, S.N., van Oers, M.M., Ros, V.I., 2019. Where the baculoviruses lead, the caterpillars follow: baculovirus-induced alterations in caterpillar behaviour. Curr. Opin. Insect Sci. 33, 30–36. https://doi.org/10.1016/j.cois.2019.02.008

Goulson, D., 1997. Wipfelkrankheit: Modification of host behaviour during baculoviral infection. Oecologia 109, 219–228. https://doi.org/10.1007/s004420050076

Granados, R.R., Lawler, K.A., 1981. In vivo pathway of Autographa californica baculovirus invasion and infection. Virology 108, 297–308. https://doi.org/10.1016/0042-6822(81)90438-4

Han, Y., van Houte, S., Drees, G.F., van Oers, M.M., Ros, V.I.D., 2015. Parasitic manipulation of host behaviour: Baculovirus SeMNPV EGT facilitates tree-top disease in spodoptera exigua larvae by extending the time to death. Insects 6, 716–731. https://doi.org/10.3390/insects6030716

Han, Y., Van Houte, S., Van Oers, M.M., Ros, V.I.D., 2018. Timely trigger of caterpillar zombie behaviour: Temporal requirements for light in baculovirus-induced tree-Top disease. Parasitology 145, 822–827. https://doi.org/10.1017/S0031182017001822

Hoover, K., Grove, M., Gardner, M., Hughes, D.P., McNeil, J., Slavicek, J., 2011. A gene for an extended phenotype. Science (80-.). 333, 1401. https://doi.org/10.1126/science.1209199

Isaac, R.E., Taylor, C.A., Hamasaka, Y., Nässel, D.R., Shirras, A.D., 2004. Proctolin in the post-genomic era: New insights and challenges. Invertebr. Neurosci. 5, 51–64. https://doi.org/10.1007/s10158-004-0029-5

Johnson, B.R., Atallah, J., Plachetzki, D.C., 2013. The importance of tissue specificity for RNA-seq: Highlighting the errors of composite structure extractions. BMC Genomics 14. https://doi.org/10.1186/1471-2164-14-586

Kamita, S.G., Nagasaka, K., Chua, J.W., Shimada, T., Mita, K., Kobayashi, M., Maeda, S., Hammock, B.D., 2005. A baculovirus-encoded protein tyrosine phosphatase gene induces enhanced locomotory activity in a lepidopteran host. Proc. Natl. Acad. Sci. 102, 2584–2589. https://doi.org/10.1073/pnas.0409457102

Katsuma, S., Koyano, Y., Kang, W., Kokusho, R., Kamita, S.G., 2012. The Baculovirus Uses a Captured Host Phosphatase to Induce Enhanced Locomotory Activity in Host Caterpillars. PLoS Pathog. 8, e1002644. https://doi.org/10.1371/journal.ppat.1002644

Kharbanda, N., Jalali, S.K., Ojha, R., Bhatnagar, R.K., 2015. Temporal expression profiling of novel spodoptera litura nucleopolyhedrovirus-encoded microRNAs upon infection of Sf21 cells. J. Gen. Virol. 96, 688–700. https://doi.org/10.1099/jgv.0.000008

Kinoshita, M., Homberg, U., 2017. Brain Evolution by Design. From Neural Origin to Cognitive Architecture-Chapter 6: Insect Brains: Minute Structures Controlling Complex Behaviors. Springer US. https://doi.org/10.1007/978-4-431-56469-0_12

Kong, M., Zuo, H., Zhu, F., Hu, Z., Chen, L., Yang, Y., Lv, P., Yao, Q., Chen, K., 2018. The interaction between baculoviruses and their insect hosts. Dev. Comp. Immunol. 83, 114–123. https://doi.org/10.1016/j.dci.2018.01.019

Lacey, L.A., Grzywacz, D., Shapiro-Ilan, D.I., Frutos, R., Brownbridge, M., Goettel, M.S., 2015. Insect pathogens as biological control agents: Back to the future. J. Invertebr. Pathol. 132, 1–41. https://doi.org/10.1016/j.jip.2015.07.009

Lange, A.B., Orchard, I., 2006. Handbook of Biologically Active Peptides - Chapter 27: Proctolin in Insects. Elsevier Inc. https://doi.org/10.1016/B978-012369442-3/50030-1

Langmead, B., Salzberg, S.L., 2012. Fast gapped-read alignment with Bowtie 2. Nat. Methods 9, 357–360. https://doi.org/10.1038/nmeth.1923

Li, B., Dewey, C.N., 2011. RSEM: Accurate transcript quantification from RNA-Seq data with or without a reference genome. BMC Bioinformatics 12. https://doi.org/10.1186/1471-2105-12-323.

Livak, K.J., Schmittgen, T.D., 2001. Analysis of relative gene expression data using real-time quantitative PCR and the 2-ΔΔCT method. Methods 25, 402–408. https://doi.org/10.1006/meth.2001.1262

Llopis-Giménez, A., Han, Y., Kim, Y., Ros, V.I.D., Herrero, S., 2019. Identification and expression analysis of the Spodoptera exigua neuropeptidome under different physiological conditions. Insect Mol. Biol. 28, 161–175. https://doi.org/10.1111/imb.12535

Martínez-Solís, M., Jakubowska, A.K., Herrero, S., 2017. Expression of the lef5 gene from Spodoptera exigua multiple nucleopolyhedrovirus contributes to the baculovirus stability in cell culture. Appl. Microbiol. Biotechnol. 101, 7579–7588. https://doi.org/10.1007/s00253-017-8495-y

Moscardi, F., Marlinda Lobo de Souza, Castro, M.E.B. de, Moscardi, M.L., Szewczyk, B., 2011. Agricultural and environmental applications - Chapter 16: Baculovirus Pesticides: Present State and Future Perspectives, Microbes and Microbial Technology. https://doi.org/10.1007/978-1-4419-7931-5

Nässel, D.R., Homberg, U., 2006. Neuropeptides in interneurons of the insect brain. Cell Tissue Res. 326, 1–24. https://doi.org/10.1007/s00441-006-0210-8

O’Reilly, D.R., Lois K. Miller, 1991. Improvement of a baculovirus pesticide by deletion of the egt gene. Biotechnology 9, 1086–1089.

O’Shea, M., Bishop, C.A., 1982. Neuropeptide Proctolin associated with an identified skeletal motoneuron. J. Neurosci. 2, 1242–1251.

Orchard, I., Belanger, J.H., Lange, A.B., 1989. Proctolin: A review with emphasis on insects. J. Neurobiol. 20, 470–496. https://doi.org/10.1002/neu.480200515

Orchard, I., Lee, D.H., da Silva, R., Lange, A.B., 2011. The proctolin gene and biological effects of proctolin in the blood-feeding bug, Rhodnius prolixus. Front. Endocrinol. (Lausanne). 2, 1–10. https://doi.org/10.3389/fendo.2011.00059

Ormerod, K.G., LePine, O.K., Bhutta, M.S., Jung, J., Tattersall, G.J., Mercier, A.J., 2016. Characterizing the physiological and behavioral roles of proctolin in Drosophila melanogaster. J. Neurophysiol. 115, 568–580. https://doi.org/10.1152/jn.00606.2015

Passarelli, A.L., 2011. Barriers To Success: How baculoviruses establish efficient systemic infections. Virology 411, 383–392. https://doi.org/10.3138/9781442622739-013

Schoofs, L., Loof, A. De, Hiel, M.B. Van, 2017. Neuropeptides as Regulators of Behavior in Insects. Annu. Rev. Entomol. https://doi.org/10.1146/annurev-ento-031616-035500

Singh, J., Singh, C.P., Bhavani, A., Nagaraju, J., 2010. Discovering microRNAs from Bombyx mori nucleopolyhedrosis virus. Virology 407, 120–128. https://doi.org/10.1016/j.virol.2010.07.033

Smirnoff, W.A., 1965. Observations on the effect of virus infection on insect behavior. J. Invertebr. Pathol. 7, 387–388. https://doi.org/10.1016/0022-2011(65)90017-0

Szewczyk, B., Hoyos-Carvajal, L., Paluszek, M., Skrzecz, I., Lobo De Souza, M., 2006. Baculoviruses - Re-emerging biopesticides. Biotechnol. Adv. 24, 143–160. https://doi.org/10.1016/j.biotechadv.2005.09.001

Tang, Q., Qiu, L., Li, G., 2019. Baculovirus-Encoded MicroRNAs: A Brief Overview and Future Prospects. Curr. Microbiol. 76, 738–743. https://doi.org/10.1007/s00284-018-1443-y

Taylor, C.A.M., Winther, Å.M.E., Siviter, R.J., Shirras, A.D., Isaac, R.E., Nässel, D.R., 2004. Identification of a Proctolin Preprohormone Gene (Proct) of Drosophila melanogaster: Expression and Predicted Prohormone Processing. J. Neurobiol. 58, 379–391. https://doi.org/10.1002/neu.10301

Torquato, E.F.B., De Miranda Neto, M.H., Brancalhão, R.M.C., 2006. Nucleopolyhedrovirus infected central nervous system cells of Bombyx mori (L.) (Lepidoptera: Bombycidae). Neotrop. Entomol. 35, 70–74. https://doi.org/10.1590/S1519-566X2006000100010

Untergasser, A., Nijveen, H., Rao, X., Bisseling, T., Geurts, R., Leunissen, J.A.M., 2007. Primer3Plus, an enhanced web interface to Primer3. Nucleic Acids Res. 35, W71–W74. https://doi.org/10.1093/nar/gkm306

Van Hiel, M.B., Van Loy, T., Poels, J., Vandersmissen, H.P., Verlinden, H., Badisco, L., Vanden Broeck, J., 2010. Advances in Experimental Medicine and Biology V. 692: Neuropeptide Systems as Targets for Parasite and Pest Control - Chapter 11: Neuropeptide Receptors as Possible Targets for Development of Insect Pest Control Agents. https://doi.org/10.1007/978-1-4419-6902-6_8

van Houte, S., Ros, V.I.D., Mastenbroek, T.G., Vendrig, N.J., Hoover, K., Spitzen, J., van Oers, M.M., 2012. Protein Tyrosine Phosphatase-Induced Hyperactivity Is a Conserved Strategy of a Subset of Baculoviruses to Manipulate Lepidopteran Host Behavior. PLoS One 7. https://doi.org/10.1371/journal.pone.0046933

Van Houte, S., Van Oers, M.M., Han, Y., Vlak, J.M., Ros, V.I.D., 2015. Baculovirus infection triggers a positive phototactic response in caterpillars to induce “tree-top” disease. Biol. Lett. 11. https://doi.org/10.1098/rsbl.2015.0132

Walker, L.E., Bloomquist, J.R., 1999. Pharmacology of contractile responses in the alimentary system of caterpillars: Implications for insecticide development and mode of action. Ann. Entomol. Soc. Am. 92, 902–908. https://doi.org/10.1093/aesa/92.6.902

Witten, J.L., O’Shea, M., 1985. Peptidergic innervation of insect skeletal muscle: Immunochemical observations. J. Comp. Neurol. 242, 93–101. https://doi.org/10.1002/cne.902420106

Wu, Q., Brown, M.R., 2006. Signaling and Function of Insulin-Like Peptides in Insects. Annu. Rev. Entomol. 51, 1–24. https://doi.org/10.1146/annurev.ento.51.110104.151011

