## Supplementary Table for "Neuropeptidome regulation after baculovirus infection. A focus on proctolin and its relevance in locomotion and digestion"

**Supplementary Table 1. Sequence list of the primers used for RT-qPCR.**

| Gene | Primer | Primer (5'-3') |
|---|---|---|
| AcMNPV proctolin | Forward | AGAACGATGCAAGTTTTCCT |
| | Reverse | GATGCATAGCATGCGGTACC |
| Adipokinetic hormone 1 | Forward | CTATTCCTGGCTTGCCTTTG |
| | Reverse | CAATTGTCTCGTCGCTTCTG |
| Allatostatin C 1 | Forward | AGAACACTCTAGTGGCGCATC |
| | Reverse | AAGTTGCAGCAGCAGTTTGC |
| Allatotropin | Forward | GACTTTGGCAAAAGGAGTGG |
| | Reverse | TCCCAGAAGTTGTCCAAACC |
| ATPsynthase subunit C | Forward | TCCTGCTGTTGTTCGCTTTC |
| | Reverse | CCACACATTCGATTCCATGGC |
| Bursicon subunit α | Forward | GCGATACCTTCTTTCGCTTG |
| | Reverse | TACAGGTCCGTTCCATTTGC |
| Bursicon subunit β | Forward | CGAAAACTACTTGCGGGAAC |
| | Reverse | TAGCACTTGCAATCGTCAGG |
| CCHamide 1 | Forward | AAGTGTTCGCGTTGCTTCTC |
| | Reverse | AAGGGTCGACGTTTTGCTAC |
| CCHamide 2 | Forward | GGCGCAAATGTTCTTAGCTG |
| | Reverse | TGATCCATGGCAGAAGTGTC |
| Corazonin | Forward | ATGGTAACCAACACGACCCTAC |
| | Reverse | ATCCGCGAGAGTATTGGAAG |
| AcMNPV DNApol | Forward | GGGTCAGGCTCCTCTTTGC |
| | Reverse | TTACGCAGCCATCACAAACAC |
| Diuretic hormone 41 | Forward | ATTCACAGACTGGCAGTGGTC |
| | Reverse | TCGGATTAACTGGGCCTTC |
| Ecdysis triggering hormone | Forward | TCCGTTCTATTGGCGTACTG |
| | Reverse | CATAGATCCTTGGTCGAACG |
| FMRFamide | Forward | GCGAGGGACCATTTCATTAG |
| | Reverse | ATGACACGCTTCCAAAGACC |
| Insuline-like peptide 1 | Forward | TTCGTCTGCTGAAGAGCATC |
| | Reverse | TGGTGATTGTGTTGGTGGTG |
| Insuline-like peptide 2 | Forward | CTGCTGATGAAGACGATTCC |
| | Reverse | CAAGCTCGTACATAGGCTTGC |
| Insuline-like precursor polypeptide Aa | Forward | TGGTTCTTGTGGTCCTCGTAG |
| | Reverse | CCAGCATCACGTTTCTCTTC |
| Insuline-like precursor polypeptide B | Forward | TGTGTTTTTCGCTGTTGTGC |
| | Reverse | ACCAGCTTCGCTTGTTGAG |
| ITG-like 1 | Forward | ATTGGTGTGCGAGGAAGTTG |
| | Reverse | TCGCTTGAAGAGTTGCAGTC |
| ITG-like 2 | Forward | TCACCTTACCGCCATTATCC |
| | Reverse | GATGCGGTTACCTTCCAAAG |
| Ion transport peptide | Forward | TTCACCCTCGAGTGCAAAG |
| | Reverse | ATAGAGCTGTGGTTCCCTGAAG |
| Leucokinin | Forward | CCGCAGTACATCAACAATGG |
| | Reverse | TTGAGGAGGGCAAGTTGAAG |
| Neuroparsin | Forward | AAGTATGCGCTAGGACGTTAGG |
| | Reverse | ACGAACCATCGAGACACTTG |
| Neuropeptide F 1 | Forward | TCCGTATGCTGCAAGAACTG |
| | Reverse | CGTCAGAACGCTTTCCAAAC |
| Neuropeptide-like precursor 1a | Forward | GAAGGACGAAGCGAATGAAG |
| | Reverse | TTTCAGACAGAGGAGCGATG |
| Orcokinin | Forward | AGTCCATACGGAAGCAAACG |
| | Reverse | TTCTTCACGAACGTGTCCAG |
| PBAN-DH | Forward | AAGGATGGCGGTTCAGATAG |
| | Reverse | AGGATCGTTTTCCGAGTCTG |
| Proctolin | Forward | GCTAACAAACCGCCAACAAC |
| | Reverse | TAATCTGGGCCTTTGTCCTG |
| Prothoracicostatic peptide 1 | Forward | GAGGCTGGAATGACATGAGC |
| | Reverse | GTTGGCCCATTTTTCAGGTC |
| Short neuropeptide F | Forward | GCCGATCGGACAACAATATG |
| | Reverse | AGCCGTTAGCTTTCACTTCC |
| SIFamide | Forward | TTATCCTGGCCTTGTGCTTC |
| | Reverse | CTCAACAACTCCCCTTTTGC |
| Sulfakinin | Forward | GATGTTGGTGCATTGCTACG |
| | Reverse | TCATCGAAGGCGTCATCAG |
| Trissin 2 | Forward | ACAGCTTCACGAGCTAATGG |
| | Reverse | TAAGGTCGGTATTCGCTTGC |
